# AAV6-HCN4t-mediated biological pacing as a potential life-saving therapy for congenital complete heart block

**DOI:** 10.1101/2025.07.31.666254

**Authors:** Jianan Wang, Evelien E. Birza, Veronique M.F. Meijborg, Arie O. Verkerk, Aina Cervera-Barea, Timo Jonker, Ruud N. Visser, Rinske Sparrius, Mischa Klerk, Arie R. Boender, Yuting Yang, Emiel J.M. Kramer, Ganghyun Kwak, Kyung Ho Park, Larry C. Park, Joyce Visser, Marlijn S. Jansen, Silke Schrödel, Mudit Mishra, Christian Thirion, Joost P.G. Sluijter, Saskia C.A. de Jager, Klaus Neef, Nico A. Blom, Monique C. Haak, Osne F. Kirzner, Hanno L Tan, Vincent M. Christoffels, Gerard J.J. Boink

**Affiliations:** Department of Medical Biology, Amsterdam Cardiovascular Sciences, Amsterdam University Medical Centers, University of Amsterdam, Meibergdreef 15, 1105 AZ, Amsterdam, the Netherlands; Experimental Cardiology Laboratory, Regenerative Medicine Center Utrecht, Circulatory Health Research Center, University Medical Center Utrecht, Utrecht University, Uppsalalaan 8, 3584 CT, Utrecht, the Netherlands; Department of Experimental Cardiology, Amsterdam Cardiovascular Sciences, Amsterdam University Medical Centers, University of Amsterdam, Meibergdreef 15, 1105 AZ, Amsterdam, the Netherlands; PacingCure BV, Roetersstraat 35, 1018 WB, Amsterdam, the Netherlands; Naason Science Inc, 123 Saengmyung-ro, 28644, Cheongju, Korea; Revvity Gene Delivery GmbH, Am Haag 6, 82166, Graefelfing, Germany; Netherlands Heart Institute, Moreelsepark 1, 3511 EP, Utrecht, the Netherlands; Division of Pediatric Cardiology, Department of Pediatrics, Willem-Alexander Children’s Hospital, Leiden University Medical Center, Albinusdreef 2, 2333 ZG, Leiden, the Netherlands; Department of Obstetrics and Fetal Medicine, Leiden University Medical Center, Albinusdreef 2, 2333 ZG, Leiden, Netherlands; Department of Anaesthesiology, Amsterdam University Medical Centers, De Boelelaan 1117, 1081 HV, Amsterdam, the Netherlands; Department of Cardiology, Amsterdam Cardiovascular Sciences, Amsterdam University Medical Centers, University of Amsterdam, Meibergdreef 9, 1105 AZ, Amsterdam, the Netherlands

**Author notes:** These authors contributed equally to this work (Jianan Wang, Evelien E. Birza).

## Abstract

Congenital complete heart block (CCHB) is a life-threatening condition in fetuses due to severe bradycardia. Maternal administration of β-adrenergic agonists is used to increase fetal heart rates, but its effectiveness is limited and lost over time most likely due to insufficient expression of HCN channels in some individuals. We report the development of an injectable gene therapy that produces reliable cardiac pacemaker function in the presence of β-adrenergic stimulation. Intramyocardial injection of adeno-associated viral serotype 6 vectors expressing HCN4t (AAV6-HCN4t) into the left ventricular apex significantly increased ectopic pacing frequency and heart rate in response to isoproterenol in rats with complete heart block, and this effect remained stable throughout the 4 weeks of follow-up. Injection of AAV6-HCN4t showed similar reliable biological pacing in complete heart block pigs. These results suggest that AAV6-HCN4t generates robust biological pacing in the presence of isoproterenol, providing the foundation for a potentially life-saving therapy for *in utero* CCHB.

## Main text

Congenital complete heart block (CCHB) is a rare condition that affects fetuses and is most often caused by maternal antibodies that damage the atrioventricular (AV) node of the heart. This may result in severe slowing of the heart rate (bradycardia)^1,2^ and lead to heart failure, multi organ failure, hydrops fetalis, and death. The overall mortality rate is around 20%, with most deaths occurring *in utero* or within the first three months postpartum.^3^ In severe CCHB, standard-of-care treatment includes maternal administration of β-adrenergic agonists in order to increase the fetal heart rate by activating HCN channels in the fetus’ ventricular conduction system. These channels play a critical role in the generation of ventricular escape rhythms, as evidenced by the strong sensitivity of these rhythms to ivabradine, a selective HCN blocker.^4^ However, in severe cases, the therapeutic effect of β-adrenergic stimulation is limited, resulting in only a modest increase in heart rate. Moreover, this effect may diminish overtime, likely due to insufficient HCN channel expression. To change this, we propose an *in utero* gene therapy (IUGT) that creates a β-adrenergic stimulation-dependent biological pacemaker, thereby increasing fetal heart rate to improve clinical outcomes and survival.^5^ To this end, we utilized adeno-associated viral serotype 6 vectors expressing a truncated version of the pacemaker channel gene HCN4 (AAV6-HCN4t),^6^ which is expected to generate biological pacing and respond to β-adrenergic stimulation.^7^

First, we assessed the function of AAV6-HCN4t using single-cell patch-clamp on ventricular cardiomyocytes isolated from 8-week-old healthy mice that were injected with AAV6-HCN4t or AAV6-nanoluciferase (AAV6-Nluc) as control (**Fig. 1a**). Voltage clamp experiments revealed that HCN4t-expressing cells generated hyperpolarization-activated time-dependent pacemaker current (*I*_f_) while control cells did not (**Fig. 1b,c**). Under non-stimulated conditions, only 2 out of 12 control cells exhibited spontaneous action potentials (APs), whereas all 10 HCN4t-expressing cells exhibited spontaneous APs (**Fig. 1d,e**). The maximal diastolic potentials (MDP) in HCN4t-expressing cells were less negative and closer to threshold than those in control cells (**Fig. 1f**). During overdrive stimulation at 4 Hz, HCN4t-expressing cells exhibited less negative MDPs and longer AP duration at 90% repolarization as compared to control cells (**Extended Data Fig. 1**), consistent with results from previous HCN4 overexpression studies.^8^ Together, these results suggest that HCN4t expression can induce spontaneous APs in cardiomyocytes through generation of *I*_f_.

**Figure 1:**
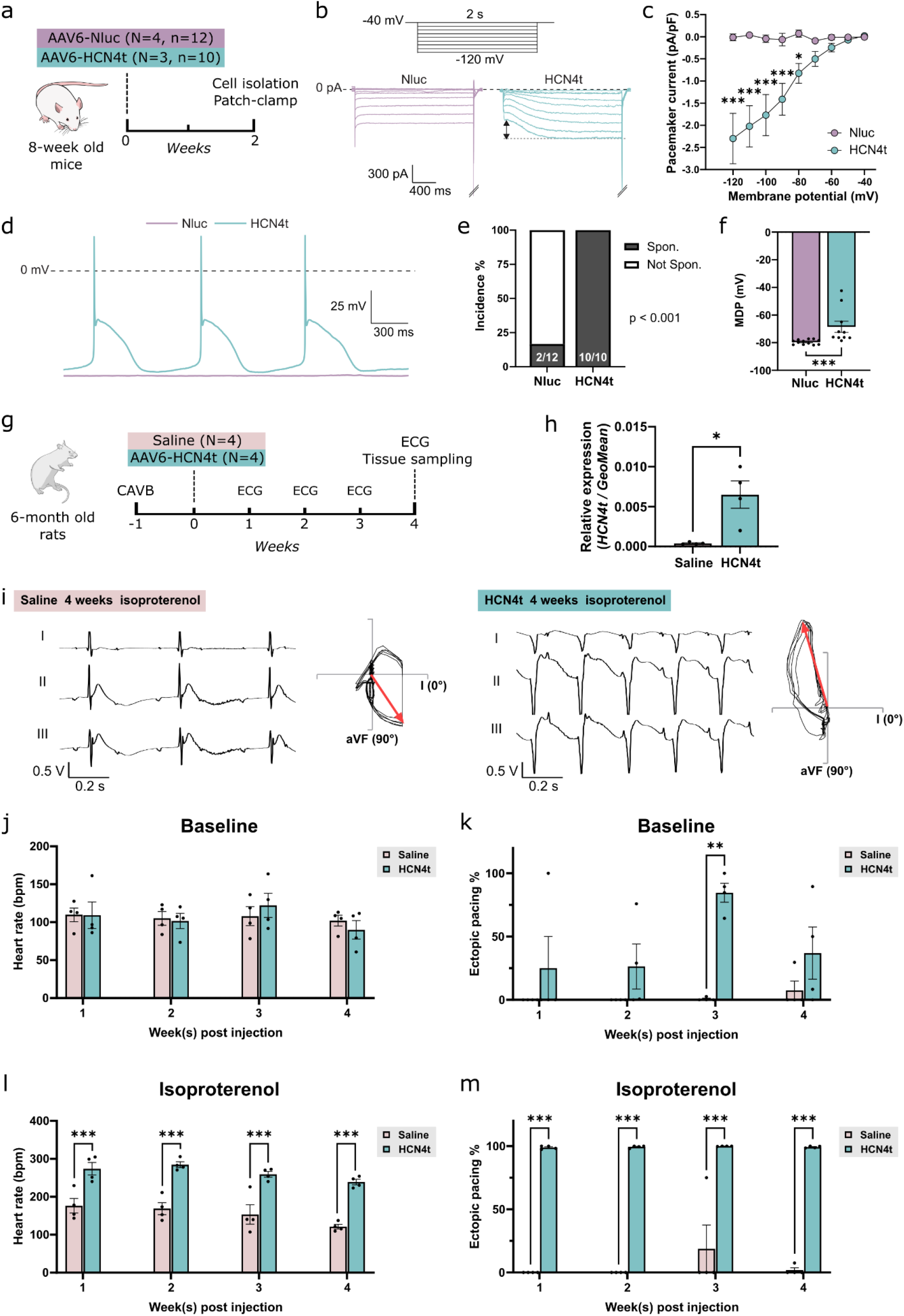
AAV6-HCN4t induces spontaneous APs in cardiomyocytes and generates biological pacing in CAVB rats. (**a**) Experimental design for the mouse single-cell electrophysiology studies. N, number of mice; n, number of cells. (**b**) Typical examples of membrane currents in a control and a HCN4t-expressing cardiomyocyte. The arrows indicate the time-dependent, hyperpolarization-activated *I*_f_. Voltage protocols used are shown at the top. (**c**) Current-voltage relationships of the hyperpolarization-activated currents. (**d**) Typical spontaneous APs (not stimulated). (**e**) Incidence of spontaneous activity. Numbers of spontaneously active and total cells are shown in the bar charts. (**f**) MDPs (not stimulated). (**g**) Experimental design for the rat studies. (**h**) Transgene expression level. (**I**) Example ECG tracings and frontal VCG plots following isoproterenol administration. Red arrows denote QRS axes. (**j**) Percentage of ectopic pacing and (**k**) average heart rates at baseline (in the absence of isoproterenol). (**l**) Percentage of ectopic pacing and (**m**) average heart rates following isoproterenol administration. Data are shown as mean ± SEM. Data were compared using a repeated measures two-way ANOVA with *post-hoc* Fisher’s least significant difference (LSD) test (c, j-m), Fisher’s exact test (e), or Mann-Whitney test (f and h). *p < 0.05; **p < 0.01; ***p < 0.001.

We then studied whether injecting AAV6-HCN4t into the heart’s left ventricle (LV) could generate a functional pacemaker in an established rat model of complete atrioventricular block (CAVB) (**Fig. 1g**).^9^ We injected 2.5 × 10^11^ vector genomes (vg) of AAV6-HCN4t or saline as control into the LV apex of 6-month-old CAVB rats. Electrocardiograms (ECG) were recorded weekly in absence and presence of isoproterenol, a β-adrenergic agonist. The rats were euthanized 4 weeks post-injection for tissue sampling. Successful transgene delivery was confirmed by qPCR (**Fig. 1h**). Typical ECG tracings and frontal vectorcardiogram (VCGs) plots at 4 weeks post-injection are shown in **Fig. 1i** and **Extended Data Fig. 2**. Following isoproterenol administration, ectopic pacing was constantly observed in AAV6-HCN4t-injected rats but rarely in control rats (**Fig. 1i**). VCGs from the saline-injected rats showed a QRS axis corresponding to AP initiation from the native conduction system, while those from AAV6-HCN4t-injected rats showed a QRS axis corresponding to AP initiation from the LV apex (**Fig. 1i**). In the absence of isoproterenol, no significant differences in heart rates were observed between the two groups (**Fig. 1j**). AAV6-HCN4t-injected rats had significantly more ectopic pacing at 3 weeks post-injection (**Fig. 1k**). Moreover, following isoproterenol administration, both ectopic pacing frequency and heart rates were significantly higher in AAV6-HCN4t-injected rats than in control rats (**Fig. 1l,m**). Notably, AAV6-HCN4t-mediated ectopic pacing remained stable throughout 4 weeks (**Fig. 1l,m**). These results indicate that AAV6-mediated HCN4t expression generates reliable biological pacing in CAVB rats, in the setting of β-adrenergic stimulation.

Subsequently, we studied the performance of AAV6-HCN4t-mediated biological pacing in 3-month-old CAVB pigs (experimental design shown in **Fig. 2a**), as the size and electrophysiological properties of pig hearts more closely resemble those of humans. An electronic pacemaker was implanted in the right ventricular (RV) apex, followed by radiofrequency ablation of the AV node.^10^ The electronic pacemaker was set to maintain a rate of 50 bpm to prevent severe bradycardia. We then performed a sternotomy and injected either AAV6-HCN4t or AAV6-control vectors into the LV apex at 3 sites with 1 × 10^12^ vg/site. At both 4 and 8 weeks post-injection, no significant differences in electronic pacemaker dependency were observed (**Fig. 2b**). Typical ECG tracings at baseline are shown in **Fig. 2c**, where electronic RV-paced beats were observed in both groups. Following isoproterenol administration, AAV6-HCN4t-injected pigs exhibited robust ectopic pacing (from the LV injection site), while control pigs rarely exhibited such ectopic beats (**Fig. 2d**). Quantitative ECG analysis revealed that at 4 weeks post-injection, AAV6-HCN4t-injected pigs had significantly higher average heart rates and a trend towards more ectopic pacing following isoproterenol administration compared to control pigs (**Fig. 2e,f**). At 8 weeks post-injection, AAV6-HCN4t-injected pigs showed significantly higher average heart rates and more ectopic pacing following isoproterenol administration (**Fig. 2g,h**). Furthermore, epicardial activation mapping confirmed that the electronic pacing originated from the RV apex while the ectopic biological pacing originated from LV apex injection site (**Fig. 2i-k**). Although AAV6-HCN4t expression on its own did not maintain a heart rate higher than 50 bpm, it generated significantly higher heart rates than control when β-adrenergic stimulation was provided. These data indicate that AAV6-HCN4t successfully generates robust biological pacing during β-adrenergic stimulation.

**Figure 2:**
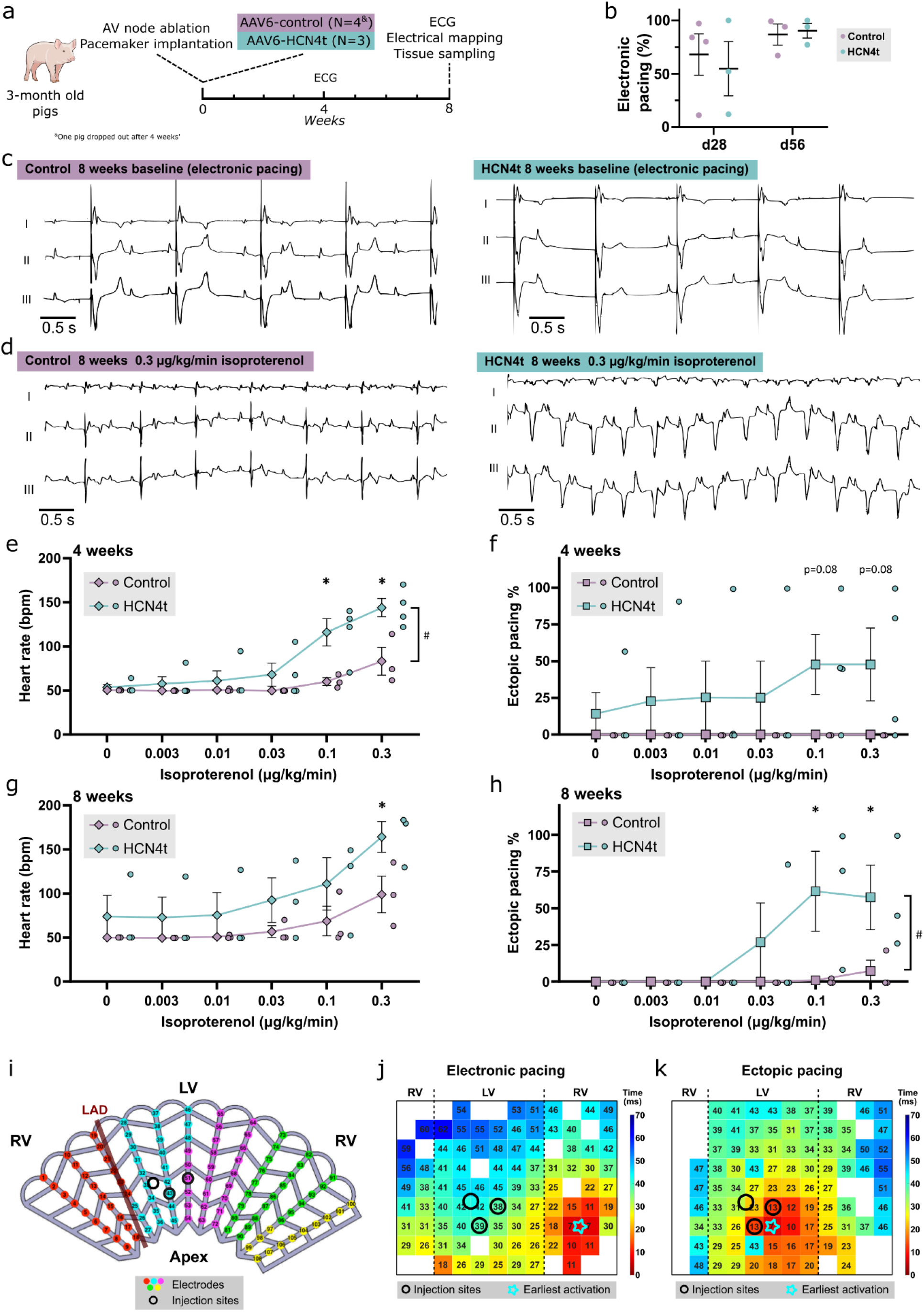
AAV-HCN4t generates biological pacing responsive to isoproterenol in CAVB pigs. (**a**) Experimental design. (**b**) Electronic pacemaker dependency at 28 and 56 days post-injection, in the absence of isoproterenol. (**c**) Example ECG tracings at baseline with electronic pacing at 50 bpm from pigs injected with AAV6-control or AAV6-HCN4t. (**d**) Example ECG tracings following infusion of 0.3 µg/kg/min isoproterenol from pigs injected with AAV6-control or AAV6-HCN4t. (**e-h**) Average heart rate and percentage of ectopic pacing with increasing doses of isoproterenol at 4 (e and f, respectively) and 8 weeks post-injection (g and h, respectively). (**i**) Schematic electrode grid with indications of LAD (maroon line) and injection sites (circles). (**j-k**) Activation maps during electronic pacing (j) and ectopic pacing (k) with indications of injection sites (circles) and the earliest activation point (star). Data are shown as mean ± SEM. Data were compared using Mann-Whitney test (b) or mixed-effect two-way ANOVA with *post-hoc* Fisher’s LSD test (e-h). *,^#^p < 0.05.

Currently, there is no effective *in utero* pacemaker therapy for CCHB. Beta-adrenergic agonists are used to increase fetal heart rates in severe cases, but the response is limited (**Extended Data Fig. 3**).^1^ In this proof-of-concept study, we demonstrate in both rodent and porcine CAVB models that a one-time injection of AAV6-HCN4t can generate a biological pacemaker that provides significantly higher heart rates following β-adrenergic stimulation. Considering that such β-adrenergic stimulation is part of the standard-of-care, we anticipate that this AAV6-HCN4t treatment can offer a potential life-saving solution for severe CCHB. To our knowledge, this is the first gene therapy-mediated biological pacemaker study with comprehensively validated efficacy over 4-8 weeks follow-up. We expect that *in utero* delivery of this therapy is feasible by injecting directly into the myocardium under ultrasound guidance (**Extended Data Fig. 4**), an approach comparable to those currently used in *in utero* valve interventions.^11^ The success of IUGT in animal studies^12^ and the emergence of *in utero* therapies in the clinical setting^13^ further support the feasibility of such an approach. Taken together, our data provide an important basis for developing an IUGT-based biological pacemaker for first-in-human testing.

## Acknowledgments

We would like to thank Ruben Coronel and Bastiaan J. Boukens for their valuable input on pig electrophysiology studies, Lisa Jansen for her assistance with tissue collection, and Jeroen Vast for his assistance with pacemaker implantation and troubleshooting. We would like to acknowledge Biotronik for providing electronic pacemakers to support our porcine studies.

