## Supplementary file for "AAV6-HCN4t-mediated biological pacing as a potential life-saving therapy for congenital complete heart block"

### Supplementary materials

#### Methods

##### Ethical approval

Mouse and pig experiments were approved and monitored by the Animal Ethics Committees of Amsterdam UMC and UMC Utrecht, and conducted in accordance with Directive 2010/63/EU of the European Parliament and with the 'Guide for the Care and Use of Laboratory Animals'. Rat experiments were carried out according to Korea Food and Drug Administration guidelines for the care and use of laboratory animals and approved by the Institutional Animal Care and Use Committee.

##### Vector production

Adeno-associated viral (AAV) vectors used in mouse experiments harbouring chicken Tnnt2 promoter-driven Nanoluciferase or N-terminal FLAG-tagged human HCN4t and p2A-GFP were produced in Amsterdam UMC as previously described.[1] AAV vectors used in rat and pig experiments harbouring chicken Tnnt2 promoter-driven non-coding multi-cloning site sequences (AAV6-control), N-terminal FLAG-tagged human HCN4t were provided by Revvity Gene Delivery (Germany).

##### Mouse single-cell patch-clamp studies

Female FVB mice of approximatly 7-8 weeks old were injected with AAV vectors at the left ventricular apex as previously described.[1] Twelve and ten cells isolated from four and three mice were used for control and HCN4t groups, respetively. Single cells were obtained by enzymatic isolation as previously described in detail.[1] Membrane potentials and currents were recorded using the amphotericin perforated patch-clamp technique with an Axopatch 200B amplifier (Molecular Devices, Sunnyvale, CA, USA). The Tyrode's solution contained (in mM): NaCl 140, KCl 5.4, $CaCl_2$ 1.8, $MgCl_2$ 1.0, glucose 5.5, HEPES 5.0, pH 7.4 (NaOH). Patch pipettes were pulled from borosilicate glass (TW100F-3, World Precision Instruments Germany Gmb) using a custom-made vertical microelectrode puller. The pipettes had tip resistances of 2–2.5 MΩ after filling with the pipette solution containing (in mM): K-gluconate 125, KCl 20, NaCl 10, amphotericin-B 0.44, HEPES 10, pH 7.3 (KOH). Cell membrane capacitance ($C_m$) was estimated by dividing the time constant of the decay of the capacitive transient in response to 5 mV hyperpolarizing voltage clamp steps from –40 mV by the series resistance. Potentials were corrected for liquid junction potential, and cell capacitance ($C_m$) and series resistance ($R_s$) were compensated for >75%. Signals were low-pass filtered (cut-off frequency 5-kHz) and digitized at 5-kHz (membrane currents and spontaneous APs), and 40-KHz (APs at 4 Hz pacing).

$I_f$ was examined as inwardly directed, time-dependent current during 2-s hyperpolarizing voltage-clamp steps from a holding potential of −40 mV (see **Fig 1b**)[2] and normalized for membrane capacitance ($C_m$). APs were measured under unpaced conditions to characterize spontaneous activity and at an overdrive stimulation of 4 Hz by 3-ms, ≈20 - 40% suprathreshold current pulses through the patch pipette. We analyzed the cycle length, maximal diastolic potential (MDP), maximum AP upstroke velocity ($V_{max}$), AP amplitude (APA), and action potential duration (APD) at 20, 50, and 90% repolarization ($APD_{20}$, $APD_{50}$, and $APD_{90}$, respectively). Parameters from 10 consecutive APs were averaged.

#### Atrioventricular (AV) node ablation and vector delivery in rats

Six-month old female Sprague Dawley rats were used for experiments and housed with a day/night light cycle of 12 hours. Control and HCN4t groups each contains four animals. Anaesthesia was induced with 5% isoflurane in 2 L/min $O_2$ in the induction chamber. Rats were shaved, intubated with a 18G catheter, and placed on a heating mat to maintain body temperature. Meloxicam (5 mg/kg) and buprenorphine SR (1 mg/kg) were delivered subcutaneously for analgesia. Anaesthesia was maintained using ventilation with 2% isoflurane in 2 L/min $O_2$. Partial right thoracotomy was performed to create an ambulatory complete heart block model, as previously reported.[3] In short, the right atrium was tied with a 4.0 silk suture to expose the fat pad, and an electrical current was delivered to the fat pad at 5W for 60 seconds. If complete AV block (CAVB) was not generated, the current delivery was repeated under the same condition. Electrocardiograms (ECG) were recorded one week after CAVB induction to confirm stable CAVB. Upon validating stable CAVB, rats were randomly selected to receive saline (n = 4) or HCN4t vectors (n = 4). A partial left thoracotomy was performed to expose the apex of the heart at the fourth intercostal space and inject 2.5 × $10^{11}$ viral genomes of AAV6 vectors, suspended in 50 µL saline, into the left ventricular apex. Control rats were injected with 50µL saline. We injected 3 mg/kg rapamycin intraperitoneally every other day to improve the transduction efficiency.[4] ECGs were recorded weekly under anesthesia for 60 minutes post AAV injection. Fifteen minutes after the start of ECG recording, isoproterenol (1mg/kg) was injected intraperitoneally. Four weeks post-injection, rats were euthanized and hearts were snap-frozen for molecular analysis.

#### Pacemaker implantation and AV node ablation in pigs

Seven 12-week-old male Topigs Norsvin pigs (Van Beek SPF Varkens, Lelystad, the Netherlands) were housed, premedicated, and anesthesised similar to what has been previously described.[5] Three days prior to AV node ablation and AAV injections, all pigs received immunosuppression consisting of cyclosporine (loading dose of 20mg/kg/day on the first day and 10mg/kg/day thereafter until termination) and prednisolone (2mg/kg/day until termination). One day prior to surgery, buprenorphine patches (Transtec, 35 μg/hour) were placed on the flank for peri-surgical analgesia. On the day of AV node ablation and AAV injections, the pigs were premedicated intramuscularly with ketamine (10 mg/kg), midazolam (0.4 mg/kg) and atropine (0.05 mg/kg). Vascular access was achieved in the ear vein, followed by anesthesia induction with thiopental (4 mg/kg). After endotracheal intubation, anesthesia was maintained with a constant-rate infusion of midazolam (0.5 mg/kg/h), sufentanil (2.5 μg/kg/h) and cis-atracurium (0.7 mg/kg/h). Pigs were mechanically ventilated with a positive pressure ventilator with $FiO_2$ 0.5, 1.5 L/min and an average frequency of 14 respirations/min under continuous capnography. We cannulated a branch of the arteria femoralis to monitor blood pressure changes during surgery and we used continuous 5-lead ECG recordings to monitor heart rate. $SPO_2$ was also monitored during the whole procedure. The pigs were given a 2-hour cyclosporine infusion of 200 mg/100 mL, as oral administration of the immunosuppressive agents was not possible because of pre-surgical fasting. Baseline 12-lead ECGs were recorded for 5 minutes after which a permanent intravenous line was placed in the external jugular vein.

To prevent severe bradycardia, a single-chamber electronic pacemaker was implanted. To this end, the ventricular lead was advanced from the external jugular vein and implanted in the apical region of the right ventricle. We deemed lead implantation successful when bipolar lead impedance was lower than 1000Ω, pacing could be achieved at a threshold amplitude of ≤ 1.8V, and intrinsic heart rhythm could be sensed with an amplitude of ≥ 10mV. After connecting the pacemaker lead to the pacemaker, we programmed the pacemaker to pace in VVI pacing mode at 50 bpm with a pulse amplitude of 3.6 V and a pulse width of 0.4 ms. Then, we created a pocket for the pacemaker on the right side of the neck and checked correct sensing, threshold amplitude and impedance values once more. Implantation of the pacemaker was completed after anchoring the lead with a suture sleeve to the intima layer of the internal jugular vein and suturing the vessel.

Subsequently, we induced CAVB through radiofrequency (RF) ablation of the AV node. To this end, we obtained vascular access by cannulating the femoral vein with an 8 French sheath using the Seldinger technique, and 100U/kg of heparin was administered in the IV line to prevent blood clotting. Then, we inserted a steerable 7 French ablation catheter with a 4 mm tip through the sheath and advanced the catheter through the inferior vena cava until it was positioned across the tricuspid valve with fluoroscopic guidance. We used the catheter to detect local electrograms showing a reliable atrial and ventricle signal. Once the His bundle spike was visible between the atrial and ventricle signals in three consecutive beats, we irreversibly damaged the His bundle by RF ablation (by delivering RF energy at a temperature of 70°C for 30 seconds). In case CAVB could not be confirmed by dissociated P waves and QRS complexes on the continuous 5-lead ECG after ablating once, we repeated the ablation procedure until dissociated P waves and QRS complexes occurred.

#### Vector delivery in pigs

After AV node ablation, we injected the pigs subepicardially with AAV6-HCN4t (N = 4) or AAV6 containing multiple cloning site (AAV6-control) (N = 3). To gain subepicardial access, we provided pre-incisional analgesia with lidocaine (10 mg/mL, 7-8 mL total), and performed a sternotomy and a full pericardiotomy. Sternal homeostasis was achieved with bone wax. Next, the heart was stabilized using a cardiac positioner (Starfish™) ensuring hemodynamic stability. One suture (6-0 Prolene) was sewn in the apical region of the LV free wall as a visual reference for the injection sites. We slowly injected 200 µL of viral particle solution ($5 \times 10^{12}$ vg/mL) thrice subepicardially using an insulin syringe (BD Micro-Fine™). The pericardium was partially closed with one suture (6-0 C-1 Prolene), to cover the injection sites. Hereafter, we inserted two drains in the pleural and pericardial cavities, which we anchored with a purse-string suture (2-0 FS-1 Vicryl) followed by the closure of the thorax (5 CCS Stainless steel). The intramuscular planes were surtured (2 CTX plus and 0 MH plus, Vicryl) followed by suturing the subcutaneous layer (2-0 Vicryl) and closing the skin by a continuous transdermal suture (3-0 PS-1 Monocryl). We administered levo-bupivacaine (1-2 mg/kg) around the incision site for additional analgesia. The chest drains were removed once exudate production was resolved. Lastly, a continuous ECG recorder (Epatch, Philips, the Netherlands) was topically placed on the left side of the chest.

#### Follow-up measurements, isoproterenol administration and epicardial mapping in pigs

We interrogated the pacemaker each week and replaced the continuous ECG recorder every 2 weeks. Four weeks after the AAV injections, we evaluated ectopic pacing with and without isoproterenol. Pigs were premedicated and anesthetized as mentioned above and five minute 12-lead ECGs were recorded. We first recorded a baseline ECG without isoproterenol stimulation, followed by progressively increasing the isoproterenol dose every 5 minutes (0.003, 0.01, 0.03, 0.1, and 0.3μg/kg/min respectively). Isoproterenol was infused through the same intravenous line system as the anesthetics. Eight weeks after AAV injections, we performed the same isoproterenol procedure in which we recorded five minute 12-lead ECGs. Then, we opened the chest through sternotomy and located the suture that was placed as a visual reference for the injection sites. We wrapped a large low density (12 × 9 electrodes) epicardial mapping grid around the heart, covering the three injections sites, and fixed it in place with towel clamps. First, we measured electrical activation patterns during electronically paced beats. Then, we measured electrical activation patterns during isoproterenol stimulation. We used the lowest isoproterenol dose that showed ectopic pacing during the closed chest measurements, or 0.3 μg/kg/min in case ectopic pacing was absent during those measurements. Lastly, the pigs were heparinized and terminated by cutting the inferior vena cava. We extracted the heart and cooled it on ice to reduce warm ischemic damage before sampling. One pig from the AAV6-HCN4t group reached its humane endpoint at 33 days post-injection. Isoproterenol stimulation has therefore been performed at its termination on 33 days post-injection and the data has been added to the 4 weeks post-injection data.

#### Electrocardiogram and vectorcardiogram analyses

For detailed electrocardiogram (ECG) analyses, we defined ectopic pacing as negative signals in lead I, II and III in ECG. Ectopic beats, electronically paced beats and other beats were quantified for the calculation of average heart rate and percentage of ectopic pacing. Fifteen minutes and five minutes of ECG recording per condition were used for analysis in rats and pigs, respectively. Rat frontal vector cardiogram (VCG) plots were made by plotting lead -aVF against lead I using 5 consecutive beats, where lead aVF was calculated using formula Lead aVF = Lead II − ½ Lead I.

#### Nucleic acid isolation, cDNA synthesis and qPCR

Rat RNA was isolated using TRIzol (MAN0001271, Invitrogen) according to the manufacturer's protocol. Isolated RNA was then treated with DNAse I, amplification grade (18068015, Invitrogen) according to the manufacturer's protocol for DNA removal. Complementary DNA library was transcribed from 500-1000 ng total RNA with oligo-dT primers (125 µmol/L) and the Superscript II Reverse Transcriptase (18064014, Invitrogen). Reverse-transcription qPCR was performed using the LightCycler 480 Real-Time PCR system (05015243001, Roche) and PowerUp SYBR green master mix (A25776, Intvitrogen). Relative expression of the transgene mRNA was normalized to the geomean of *Gapdh*, *18S rRNA* and *Actb*.

#### Statistical analysis

GraphPad Prism version 10 software (GraphPad Software Inc.) was used for statistical analysis. Numerical data are presented as mean ± standard error of the mean (SEM) of at least three biological replicates. Two independent groups were compared by unpaired Mann-Whitney test and multiple independent groups were compared by analysis of variance (ANOVA) followed by *post-hoc* least significant difference test for multiple comparisons. Categorical data were presented as absolute values and percentages. Categorical data were compared by Fisher's exact test. All tests were performed two-sided and statistical significance was considered for $p < 0.05$.

### Extended data

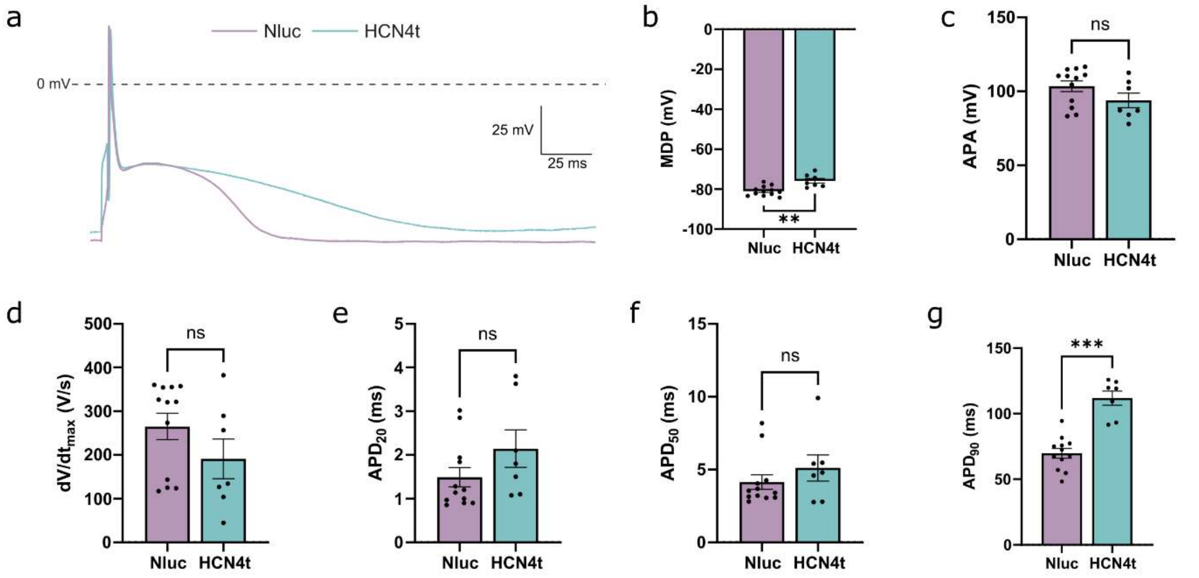


**Extended Data Fig. 1 AAV-HCN4t depolarizes MDP and prolongs AP duration in mouse cardiomyocytes.**

(**a**) Typical APs with 4 Hz overdrive stimulation. (**b**) maximal diastolic potential (MDP). (**c**) AP amplitude. (**d**) Maximal AP upstroke velocity. (**e**) AP duration at 20% repolarization. (**f**) AP duration at 50% repolarization. (**g**) AP duration at 90% repolarization. Data are shown as mean ± SEM. Data were compared using Mann–Whitney test. **p < 0.01; ***p < 0.001; ns, not significant.

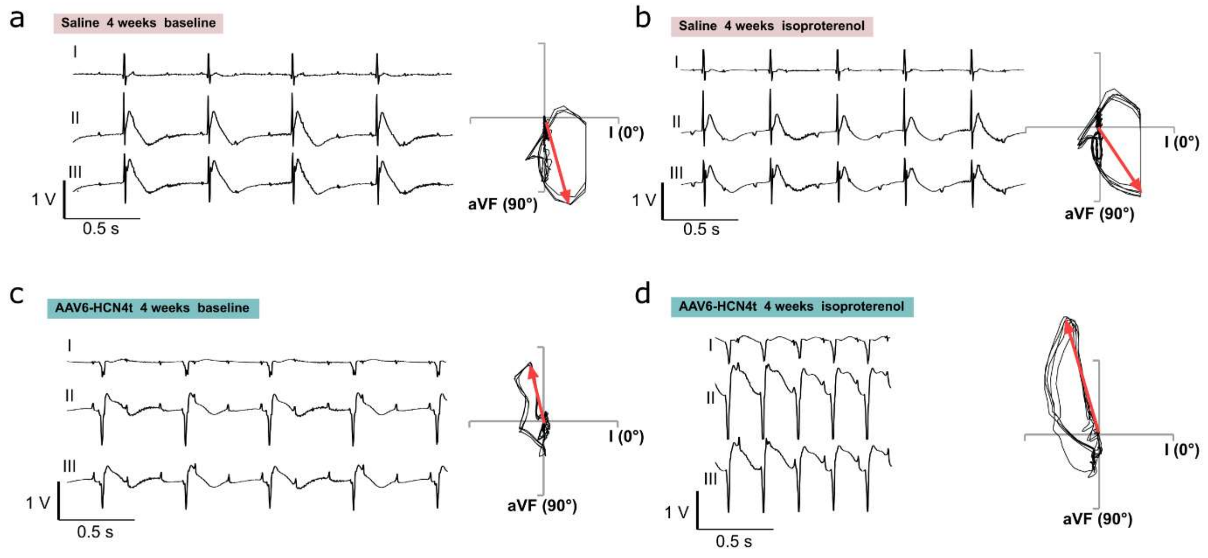


**Extended Data Fig. 2 Three-lead ECG tracings and frontal VCG measured prior to and following isoproterenol administration in CAVB rats.**

(**a**) Saline rat prior to isoproterenol admonistration. (**b**) Saline rat following isoproterenol administration. (**c**) AAV6-HCN4t prior to isoproterenol admonistration. (**d**) AAV6-HCN4t rat following isoproterenol administration. Red arrows denote QRS axes.

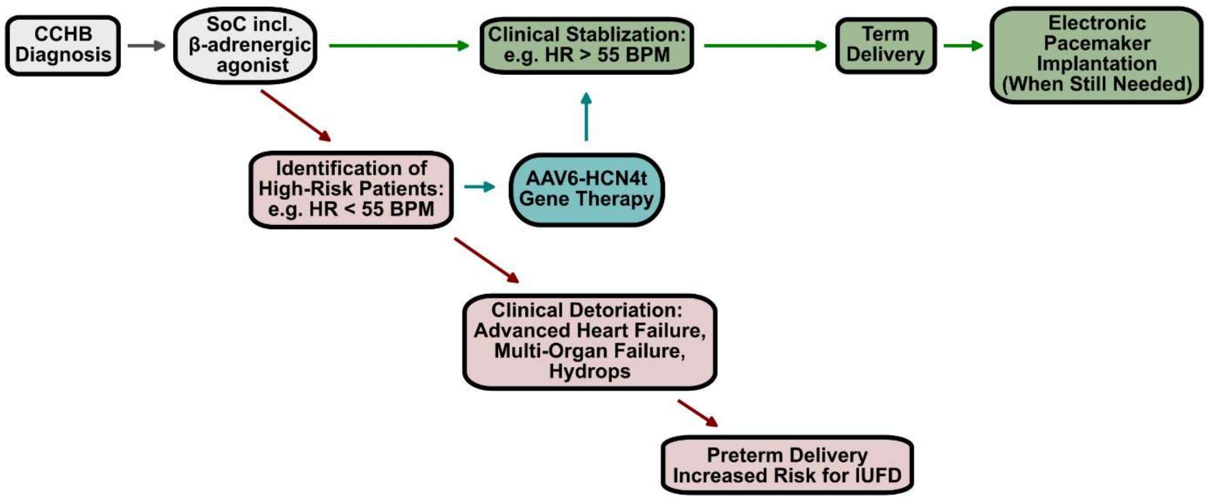


**Extended Data Fig. 3 Schematic timeline of the treatment plan for CCHB**

Once a fetus develops CCHB with a significant drop in heart rate, the standard of care including maternal corticosteroid therapy and β-adrenergic agonist are initiated to mitigate further immune-mediated myocardial damage and to increase heart rates. If the fetal heart rate remains at a non-critical level, the condition may stabilize. However, in case where the clinical situation declines (indicated by a fetal heart rate below 55 bpm), there are currently no rescue treatments available. In this high-risk context, we propose to apply AAV6-HCN4t gene therapy to stabilize the fetal heart rates and improve clinical outcomes. CCHB: Congenital complete heart Block. SoC: Standard of care. HR: Heart rate. BPM: Beats per minute IUFD: Intra-uterine fetal death.

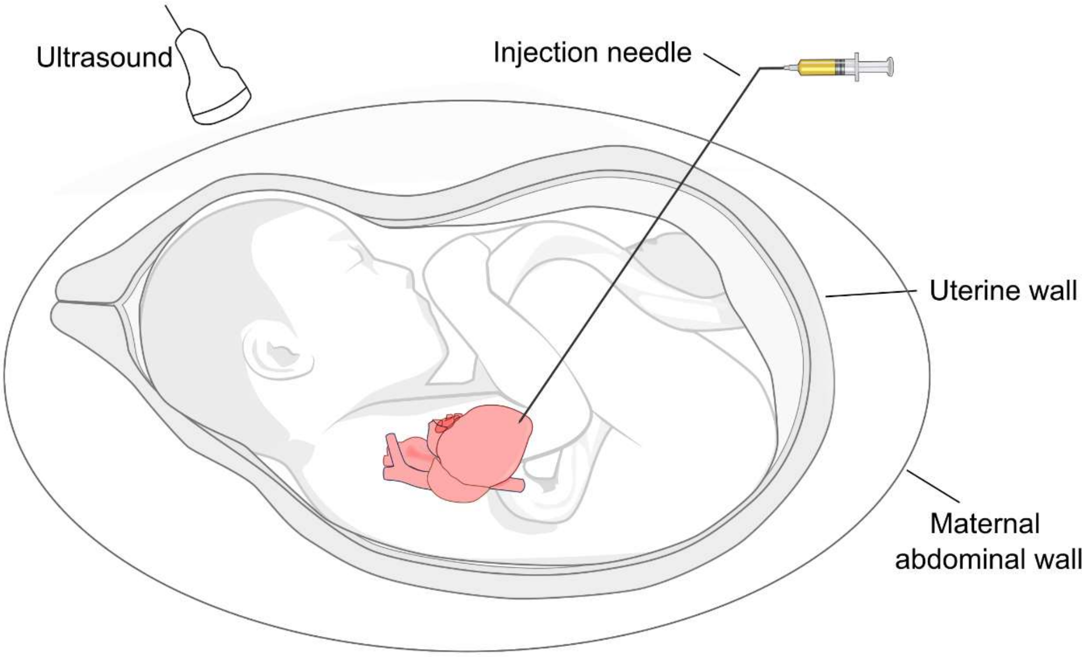


**Extended Data Fig. 4 Scheme of ultrasound-guided *in utero* intramyocardial injection**

With the use of ultrasound guidance, a needle can be inserted through the mother's abdominal wall into the amniotic cavity. The needle is advanced through the fetal chest and into the apex of the heart. Through this needle, AAV vectors are slowly injected to the targeted site.
